# Antifungal benzimidazoles disrupt vasculature by targeting one of nine β-tubulins

**DOI:** 10.1101/2020.09.15.298828

**Authors:** Riddhiman K. Garge, Hye Ji Cha, Chanjae Lee, Jimmy D. Gollihar, Aashiq H. Kachroo, John B. Wallingford, Edward M. Marcotte

## Abstract

Thiabendazole (TBZ) is an FDA-approved benzimidazole widely used for its antifungal and antihelminthic properties. We showed previously that TBZ is also a potent vascular disrupting agent and inhibits angiogenesis at the tissue level by dissociating vascular endothelial cells in newly formed blood vessels. Here, we uncover TBZ’s molecular target and mechanism of action. Using human cell culture, molecular modeling, and humanized yeast, we find that TBZ selectively targets only 1 of 9 human β-tubulin isotypes (TUBB8) to specifically disrupt endothelial cell microtubules. By leveraging epidemiological pesticide resistance data and mining chemical features of commercially used benzimidazoles, we discover that a broader class of benzimidazole compounds, in extensive use for 50 years, also potently disrupt immature blood vessels and inhibit angiogenesis. Thus, besides identifying the molecular mechanism of benzimidazole-mediated vascular disruption, this study presents evidence relevant to the widespread use of these compounds while offering potential new clinical applications.

## INTRODUCTION

The vascular system is built by the combination of *de novo* formation of blood vessels by vasculogenesis and the sprouting of new vessels from existing vessels via angiogenesis^1,2^. Imbalances in angiogenesis underlie a variety of physiological and pathological defects, including ischemic, inflammatory, and immune disorders^1,3,4^. Indeed, angiogenesis is central to tumor malignancy and cancer progression, as new blood vessels must be established to supply oxygen and nutrients to the growing tumor. Accordingly, inhibition of angiogenesis is now a well-recognized therapeutic avenue^1–5^. Defined angiogenesis inhibitors such as Avastin (FDA approved since 2004) are now in wide use in the clinic and, over the past 30 years, several dozen drugs have been approved or entered clinical trials as angiogenesis inhibitors^5–9^.

In recent years, a new class of anti-vascular drugs, termed vascular disrupting agents (VDAs), have gained attention as potential alternative therapeutics operating by distinct mechanisms^10–13^. Unlike angiogenesis inhibitors which selectively prevent the formation of new blood vessels, VDAs function by dismantling existing vasculature, making them potentially effective for therapies beyond cancer, for example in the treatment or control of macular degeneration and diabetic retinopathies^14,15^. While several VDAs have shown therapeutic potential, none have yet been approved, with several candidates still in clinical trials^11,16,17^.

Given the lengthy approval process, the failure of many drugs to succeed in clinical trials, and the high costs involved with developing new compounds, drug repurposing offers an attractive alternative for developing new therapies more quickly. We recently developed strategies to exploit data from diverse model organisms to identify both deeply conserved genetic networks as well as small molecules that may manipulate them^18–20^. This effort identified thiabendazole (TBZ) as both a novel angiogenesis inhibitor and VDA^20^.

TBZ is one of a large class of biologically active benzimidazole compounds that are widely used commercially or clinically, with applications ranging from photographic emulsions and circuit board manufacturing, to serving as one of the most common heterocyclic ring systems used for small molecule drugs^21^. The FDA approved TBZ in 1967 for human use for treating systemic fungal and helminthic infections, but it is more widely used in veterinary settings and in agricultural pesticides and preservatives. However, we found that TBZ also possesses potent vascular disrupting ability, demonstrated *in vitro* in human cell culture and *in vivo* in mice and frogs, including for retarding tumor growth and reducing intratumoral vessel density in preclinical murine xenograft models^20^.

Several other VDAs have been reported to collapse the vasculature by inhibiting microtubule polymerization dynamics *via* binding β-tubulin^11,16^. Indeed, though the basis for TBZ’s vascular disrupting action is unknown, it is proposed that TBZ’s fungicidal action is mediated *via* disrupting fungal microtubule assembly and dynamics^22,23^. In particular, mutations in β-tubulin have been frequently found to confer resistance to TBZ in parasitic/invasive fungal and nematode species^21–^ 3. However, in humans, TBZ does not generally disrupt cell growth, and even in human umbilical vein endothelial (HUVEC) cells, while it somewhat reduced tubulin protein abundance it did not elicit gross defects in the microtubule cytoskeleton^20^. At angiogenesis-inhibiting doses, the overall development of TBZ treated animals is normal^18^, consistent with TBZ’s safety record in humans and veterinary settings^24^. Therefore, we hypothesized that only certain types of human cells, such as subsets of endothelial cells involved in forming the vasculature, might be uniquely susceptible to TBZ.

Here, we experimentally determined TBZ’s specific molecular target and cellular mechanism of vascular disrupting activity. We find that TBZ disrupts microtubule growth, with increased potency in endothelial cells. Using predictive molecular modeling, human cell culture, and humanized yeast, we find TBZ predominantly targets only one of nine human β-tubulins, suggesting an explanation for its cell-type specificity. Finally, based on epidemiological data mining and chemical structures, we discovered that a larger family of benzimidazoles—in clinical and commercial use for >50 years—all act as VDAs, disrupting the vasculature in a vertebrate animal model. These newly discovered VDAs include two World Health Organization (WHO) antihelminthics (albendazole and mebendazole) administered for the treatment of human intestinal infections, one broad-spectrum antifungal/antihelminthic (fenbendazole) used to treat farm animal infections, and two banned pesticides (benomyl and carbendazim) used to prevent wild fungal and nematode mediated crop destruction. Knowledge of their vascular disrupting activities should thus inform their use in at-risk individuals (such as during pregnancy) and opens new clinical applications for these compounds.

## RESULTS

### Thiabendazole disrupts microtubule plus ends in endothelial cells

Thiabendazole exhibits broad-spectrum activity against fungal and nematode crop pests^25^, but prior to demonstration of its VDA activity, it was generally thought to lack activity in tetrapods^24^. Its binding target and mechanism of vascular action remains poorly understood^26^, although *in vitro* studies have suggested various benzimidazole compounds inhibit cell growth by interfering with microtubule polymerization^23,26–28^. Benzimidazole suppressor screens in both *Saccharomyces cerevisiae* and *Caenorhabditis elegans* have independently identified resistance mutations occuring in β-tubulin genes, giving some insight into the binding site^29,30^. The case for a β-tubulin binding site is strengthened by numerous animal and agricultural studies also demonstrating resistance mutations arising repeatedly and independently across multiple parasitic nematode and fungal species infecting farm livestock and crops^22,27,31–43^.

To examine the effects of the microtubule cytoskeleton and dynamics in the presence of TBZ, we examined the localization of GFP-tagged EB1, which labels growing microtubule plus ends and provides a proxy for microtubule dynamics in both endothelial (HUVEC) and non-endothelial (NIH3T3) human cells. Despite the grossly normal architecture of microtubules in TBZ-treated HUVECs, TBZ significantly reduced the accumulation of EB1 at microtubule plus ends (**Fig. 1A’, B’**) as compared to its control (**Fig. 1A, B**). Importantly, and consistent with the overall normal morphology and patterning of TBZ-treated embryos^20^, we found that TBZ had a substantially less robust effect on EB1 accumulation at microtubule plus ends in fibroblasts as compared to endothelial cells (**Fig. 1C-E**). These data are consistent with, and provide new insights into, previous studies showing that TBZ’s interaction with tubulin interferes with microtubule polymerization in nematodes and fungi^44,45^.

**Figure 1.**
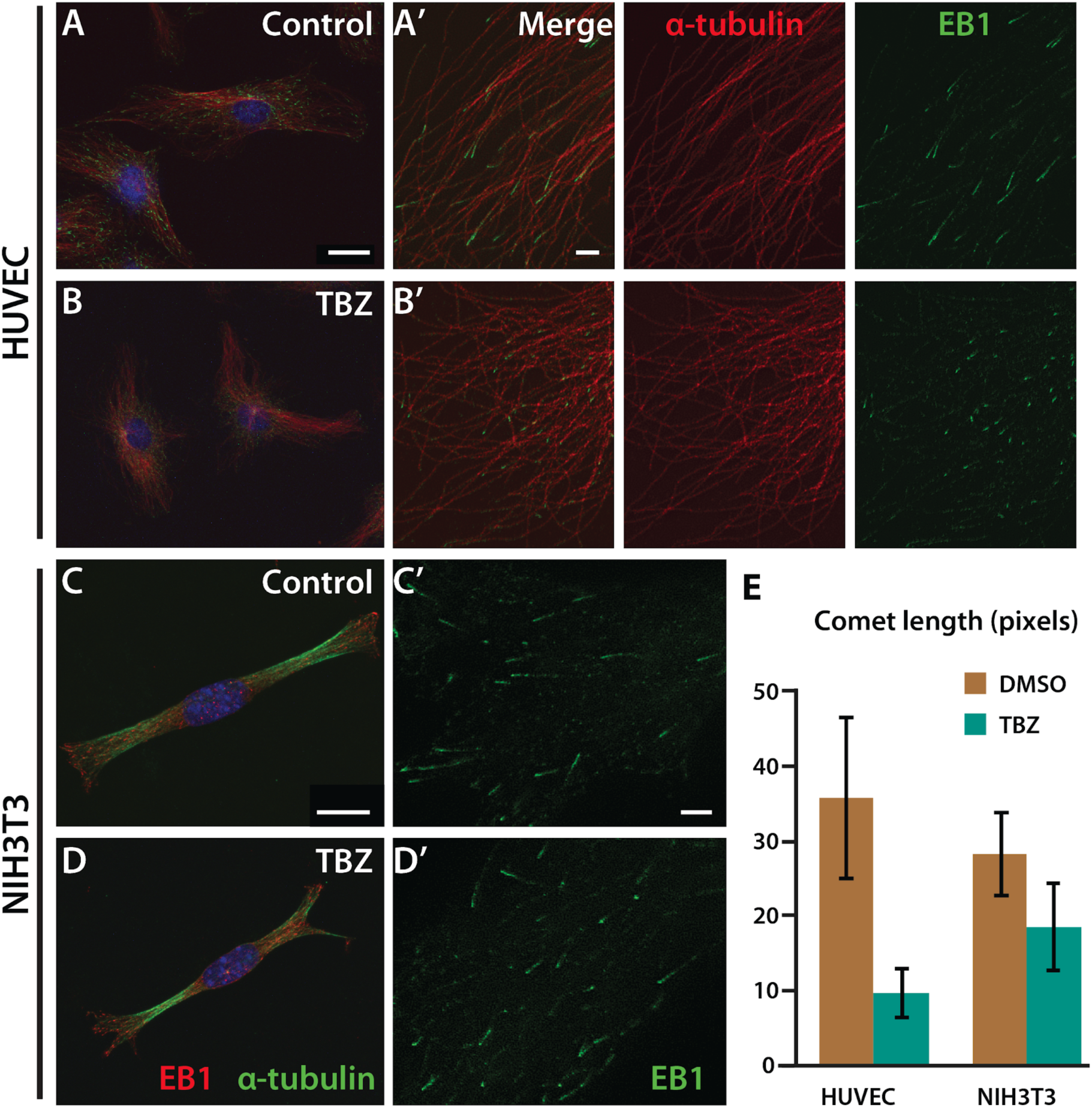
Thiabendazole (TBZ) significantly reduces EB1 comet length at microtubule plus ends in cultured human cells. Immunohistochemical analysis of α-tubulin in two human cell lines using confocal microscopy does not show a definite distinction between 1% DMSO-treated control (**A, C**) and 1% DMSO, 250 µM TBZ-treated cell lines (**B, D**), but images from super-resolution microscopy reveal that the accumulation of end-binding (EB) protein 1 at the plus end of microtubules is significantly reduced with TBZ treatment (**B**) compared to the control (**A**) in HUVECs. In NIH-3T3 cells, the reduced EB1 comet length following TBZ treatment (**D**) compared to control (**C**) is not as pronounced as in HUVECs, as quantified by comet length (**E**). Scale bars, 20 µm in (A) and (C), 2 µm in (A’) and (C’).

### Thiabendazole selectively targets TUBB8 among human β-tubulins

Three commonly observed mutations in fungal and nematode β-tubulins (F200Y, E198A, and F167Y) confer resistance to TBZ (**Fig. 2A, B**), suggesting that its binding site is in the vicinity of these residues^22,27,31–43^(**File S1**). Based on the previously observed benzimidazole suppressor mutations, we used 3D structural modeling to evaluate TBZ’s potential binding sites in a fungal β-tubulin. We first constructed 3D homology models of the *Schizosaccharomyces pombe* (fission yeast) wild-type and TBZ-resistant F200Y β-tubulins, based on the previously determined *Ovis aries* β-tubulin crystal structures (PDB: 3UT5^46^ and 3N2G^47^) as templates. We computationally refined the structures and then evaluated potential binding modes of TBZ, as detailed in the Methods, using computational docking algorithms to localize TBZ’s potential binding sites within the fungal β-tubulin structures (**Fig. S1**). We identified a binding site around F200 to be the most probable (**File S2**). We found that the preferred binding conformations of TBZ in both models (**Fig. S1**) situated close to (but distinct from) the colchicine binding site. These observations were in strong agreement with computational predictions made on parasitic β-tubulins binding benzimidazoles^27^ and recent crystal structures of other benzimidazole derivatives binding to bovine brain β-tubulins^48^.

**Figure 2.**
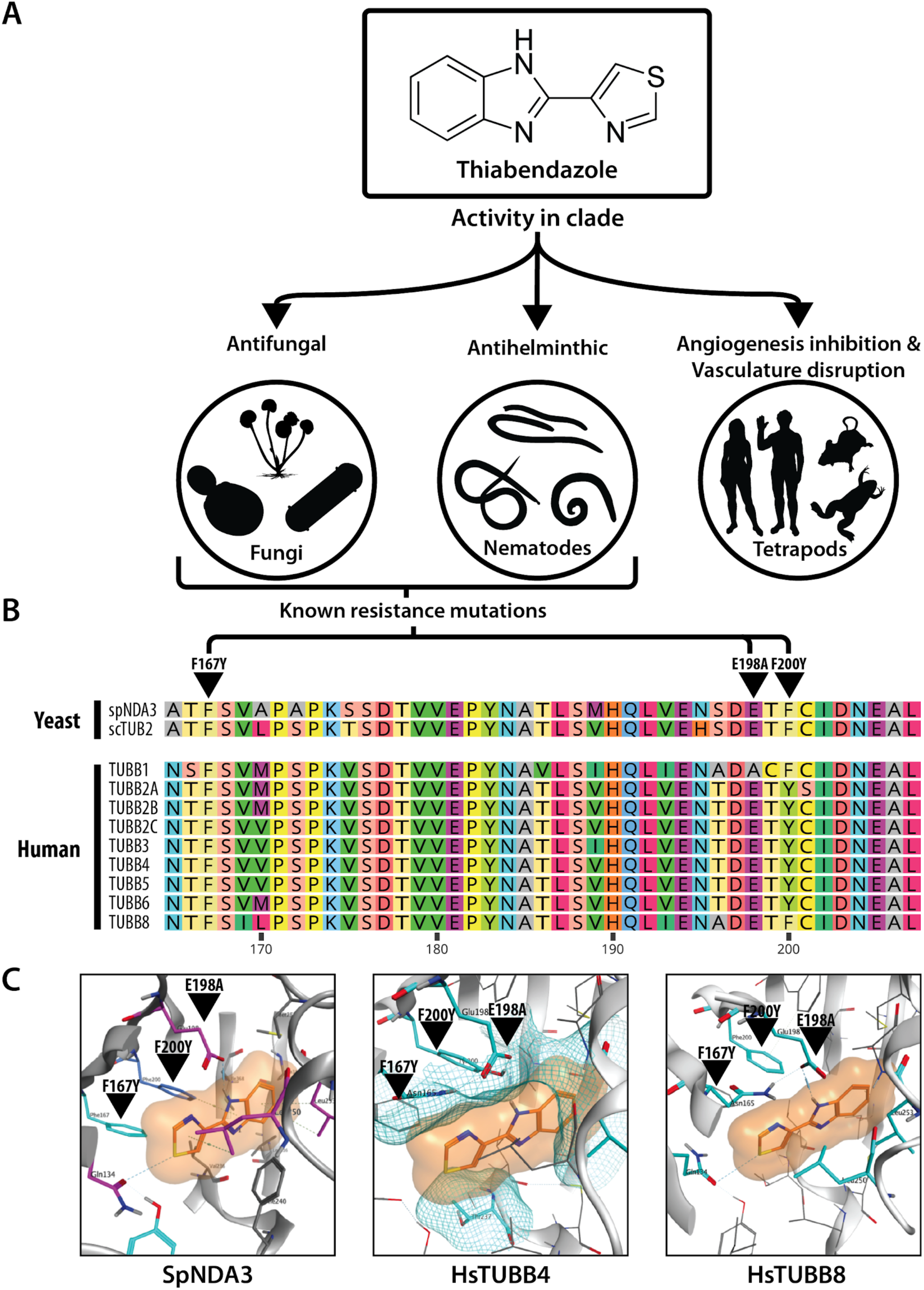
Uncovering the molecular mechanism of thiabendazole. (**A**) TBZ elicits varying activity across different clades of life being toxic to fungal and nematode clades but behaves as a vascular disrupting agent in tetrapods. (**B**) Of the 9 human β-tubulins, 8 have amino acids at positions 167, 198, and 200 that confer TBZ resistance to fungal tubulins (see Table S1), as seen in a multiple sequence alignment of human and *Schizosaccharomyces pombe* β-tubulins; only TUBB8 lacks resistance mutations. (**C**) *In silico* docking of TBZ (orange) into a homology modeled yeast β-tubulin 3D structure (see Methods) indicates TBZ is well-accommodated by a binding pocket in wild-type yeast *NDA3* that abuts the 3 major β-tubulin TBZ resistance mutation sites. In contrast, docking of TBZ into homology models of human TUBB4 and TUBB8 indicates the potential for differential binding, with TUBB8 accommodating TBZ whereas, in the case of TUBB4, TBZ is sterically blocked. Polar contacts are illustrated *via* dashed lines, and residues lining the proposed binding pocket are shown in cyan. Intramolecular hydrogen bonding between E198 and Y200 in TUBB4 reorganizes the geometry of the binding pocket. Residues involved in steric clashing are depicted with a partial mesh surface. (Note that due to steric clashes between TBZ and TUBB4 at the proposed binding pocket, TBZ was superimposed from our binding model to measure interactions).

On measuring the polar contacts and clashing energies of TBZ with tubulin, we found that the wild-type β-tubulin bound to TBZ more favorably with contact energy (−9.9 kxcal/mol) as compared to its F200Y counterpart, which showed unfavorable repulsions (+27.6 kcal/mol)(**File S2, S3**). For the wild-type protein, TBZ’s polar contacts included E198 and Q134 (**Fig. 2C, S1**). Arene-hydrogen interactions between the drug and protein included contributions from F200, L250, and L253. However, for our F200Y mutant, repulsion was observed in our fixed ligand experiments predominantly caused by unfavorable contacts made with Y200, F240, L250, and L253 (**Fig. S1**). Our analyses suggest F200Y likely forms a hydrogen bond to E198 in the TBZ-resistant mutant, thus constricting the pocket and occluding binding.

Unlike fungi, tetrapods have multiple β-tubulin isotypes (here, we use the term isotype to denote the protein products of paralogous genes, in accordance with prior tubulin literature), and their expression varies in different cells and tissue types. For example, human tubulin βI (TUBB/TUBB5) is constitutively expressed in many cells and tissues, whereas βIII (TUBB3) is exclusively enriched in neurons and the brain^49,50^. The specific roles of different β-tubulin isotypes are not yet fully understood, but recent studies indicate that their sequence diversity modulates binding affinity to tubulin-binding drugs and influences microtubule dynamics through distinct interactions with molecular motors^50,51^. The recurrence of TBZ resistance mutations at the same three loci across diverse fungi and nematodes (**File S1**) led us to hypothesize that human β-tubulin isotypes might have differential sensitivities to TBZ by virtue of incorporating resistant residues at positions 167, 198, and 200, potentially explaining both its tissue-specific effects and generally low toxicity in humans.

Indeed, multiple sequence alignment of human and yeast β-tubulin genes indicated that while F167 remained conserved across all the human isotypes, positions 198 and 200 were variable (**Fig. 2B**). Moreover, all human β-tubulin isotypes except TUBB1 and TUBB8 contain the F200Y resistance mutation. Because TUBB1 also harbors the other commonly observed E198A suppressor (**Fig. 2B**), TUBB8 is the only human β-tubulin isotype predicted by sequence to be TBZ-sensitive. Given this variability across isotypes, we next asked how the E198A and F200Y mutations would be expected to affect TBZ’s ability to bind at its predicted site in human isotypes.

We first evaluated this hypothesis computationally, by constructing 3D homology models for each of the human β-tubulin isotypes in the same manner as for the fungal model (see Methods). We then performed induced-fit docking with TBZ across our human β-tubulin models. Using a TBZ-wild-type fungal β-tubulin complex as a template, we docked TBZ into the same pocket in each of the human isotypes and measured protein-ligand interactions in the superimposed structures. In agreement with our primary sequence based predictions, TBZ fit well into the predicted binding pocket of only TUBB1 and TUBB8, which lack the F200Y mutation. Both showed favorable binding energies of −1.8 and −8.3 kcal/mol, respectively (**File S2**). The large difference in contact energy among these isotypes could be explained by position 198. In TUBB1, alanine occupies position 198, whereas TUBB8 has glutamate, which contributed heavily to the binding energy in all of our simulations when both F200 and E198 were present. Our data suggest that TBZ binding is stabilized by hydrogen bonds with residues Q134 and E198 in the presence of F200 (**Fig. S1, S2**). Taken together, our *in silico* studies predicted that TBZ should strongly bind TUBB8 and weakly bind TUBB1, but should not bind any other human β-tubulin isotypes.

### Functional assays in human endothelial cells and humanized yeast confirm TBZ specificity to human TUBB8

Given TBZ’s effects on human vascular endothelial cells and *in vivo* vascular disruption in *Xenopus* embryos^20^, we wished to test directly if TUBB8-specific binding could explain the compound’s effects. We thus asked whether resistance to TBZ could be acquired by simply supplying human β-tubulin isotypes predicted to be resistant. We tested this by two independent assays: (i) by overexpressing specific sensitive or resistant human β-tubulin isotypes in human endothelial cells and (ii) by humanizing Baker’s yeast’s β-tubulin *TUB2* to enable assays of individual human β-tubulin isotypes.

To test if microtubule dynamics in human cells could be significantly restored by supplying resistant β-tubulin isotypes, we singly transfected HUVEC cells with plasmids overexpressing either TUBB4 or TUBB8 and assayed microtubule dynamics by measuring the comet lengths of end-binding protein EB3 (**Fig. 3A**). Compared to untransfected HUVECs, we saw that overexpressing TUBB4 significantly rescued the decrease in comet length observed in TBZ-treated cells (**Fig. 3B**). Transfection with *TUBB8*, by contrast, had no effect (**Fig. 3B**). The differences became very significant after 30 minutes of exposure (**Fig. 3B**).

**Figure 3.**
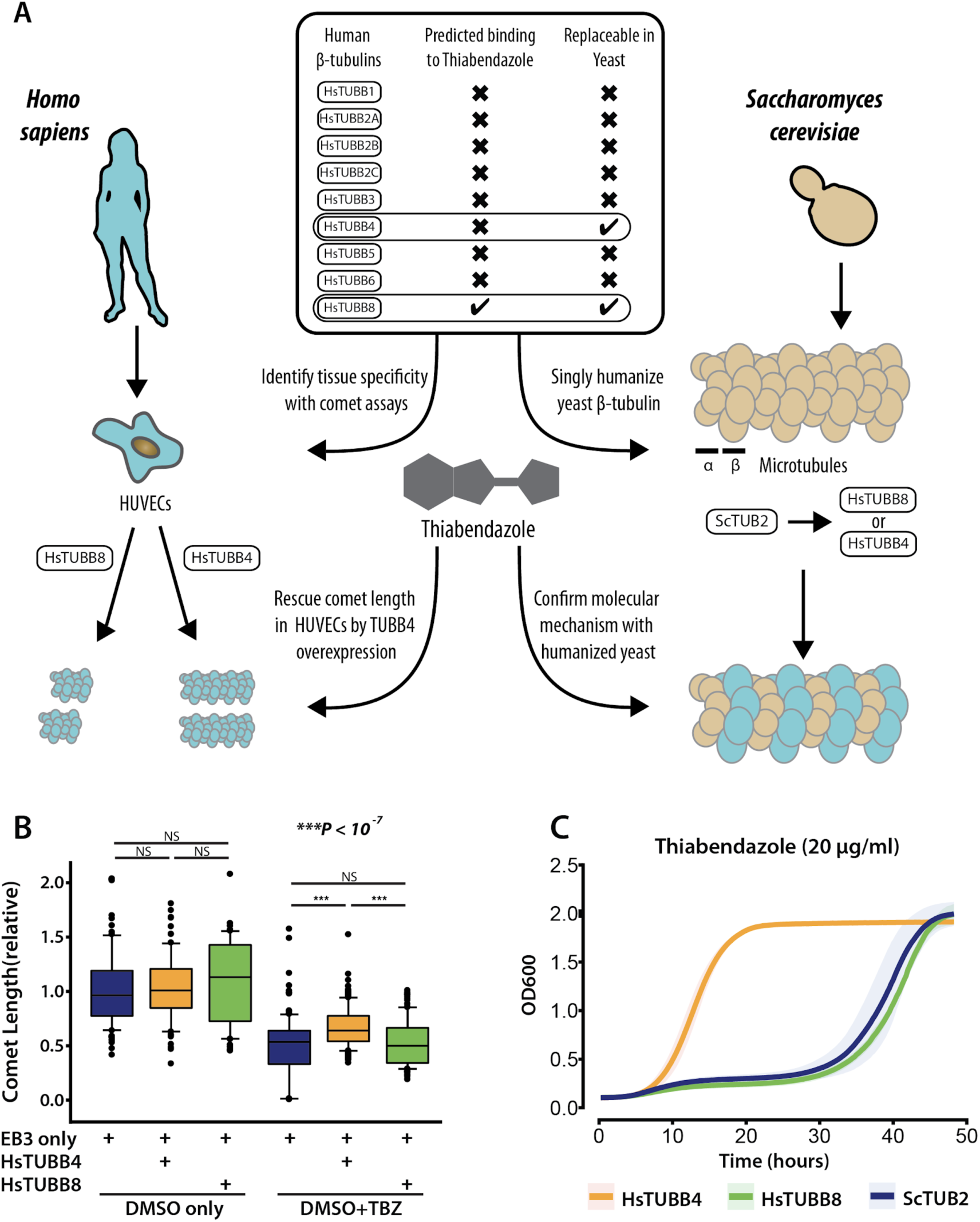
TBZ specifically inhibits the human β-tubulin TUBB8, not TUBB4, in humanized yeast and HUVEC cell culture. (**A**) Overview. TBZ’s isotype specificity was identified in 2 ways. (Left) Recombinant human β-tubulins TUBB4 and TUBB8 were individually overexpressed in HUVEC cell culture to monitor comet lengths in the presence of TBZ. (Right) Using humanized yeast wherein yeast *TUB2* was singly humanized by either of 2 replaceable human β-tubulins TUBB4 or TUBB8 to screen for differential sensitivity towards TBZ. (**B**) Reduced EB3 comet length after 1% DMSO, 250 µM TBZ treatment compared to 1% DMSO treated control. **A**. Comet length is similar in EB3, TUBB8 transfected HUVECs compared to EB3 transfected controls expressing native tubulins, but comets are longer in most EB3, TUBB4 transfected cells. **B**. Comet length is statistically similar between cells treated with 1% DMSO; however, following 30 minutes of 1% DMSO, 250 µM TBZ treatment *TUBB4* transfected cells have significantly longer EB3 comets than HUVECs with TUBB8 or expressing native tubulins. (**C**) Growth profiles of humanized yeast strains show TBZ’s isotype specificity to TUBB8. When grown in the presence of TBZ, Strains carrying the wild-type *TUB2* (blue) and human TUBB8 (green) genes are sensitive to TBZ while humanized TUBB4 strains (orange) are resistant. Mean +/-standard deviation indicated by solid lines and shaded boundaries, respectively.

As an independent assay of TBZ action on human tubulins, we turned to humanized yeast, as our previous work showed that of the nine human β-tubulins, only TUBB4 and TUBB8 could functionally replace *TUB2* in *Saccharomyces cerevisiae*^52^. From our modeling and docking data, we hypothesized that yeast strains humanized with TUBB8 would be susceptible to TBZ while humanizing with TUBB4 would confer TBZ resistance. *Saccharomyces cerevisiae* possesses 2 α-tubulins (*TUB1* and *TUB3*) that interact with *TUB2* to form tubulin heterodimers, which in turn oligomerize to form microtubules. Wild-type BY4741 haploid strains are TBZ-resistant. However, previous studies have shown that on deleting *TUB3*, yeast strains become susceptible to benzimidazoles^53^ likely due to reduced overall α-tubulin stoichiometry or possibly by TBZ occluding *TUB2*’s dimerization with *TUB1* but not *TUB3*. Therefore, we performed all our yeast replacement assays in a *tub3Δ* background, which yielded a clear growth defect in the presence of TBZ (**Fig. S3**). In order to test the effect of TBZ on human β-tubulin isotypes TUBB4 and TUBB8, we used CRISPR/Cas9 to construct yeast strains with these human isotypes in place of the endogenous *TUB2* and tested them in the presence or the absence of the drug (**Fig. 3A**). We found that strains possessing wild-type *TUB2* and human TUBB8 exhibited slow growth in the presence of TBZ (at conc. 20 µg/ml). By contrast, the strain humanized with TUBB4, which is predicted to be resistant to TBZ, grew normally in the presence of TBZ (**Fig. 3C, S3A**).

Together with our *in silico* docking data, our results in HUVECs and humanized yeast indicate that TUBB8 is uniquely TBZ-sensitive, suggesting in turn that vascular endothelial cells are selectively sensitive to its loss.

### Benzimidazole resistance patterns and chemical similarities suggest additional VDAs

Given the plethora of fungal and nematode studies on benzimidazole pesticide resistance in agriculture^22,33–36,38,40–44,54–71^ (**Fig. 4A**), we reasoned that TBZ’s molecular mechanism may extend to other commercially used benzimidazole compounds. Indeed, based on our experiments, a simple epidemiological signature should be sufficient to identify other pesticides that likely to function as vascular disrupting agents and angiogenesis inhibitors: (i) the compounds should be selectively toxic to fungal and nematode clades but demonstrate low toxicity in tetrapods, and (ii) sensitive species should specifically gain benzimidazole resistance from F167Y, E198A or F200Y β-tubulin mutations.

**Figure 4.**
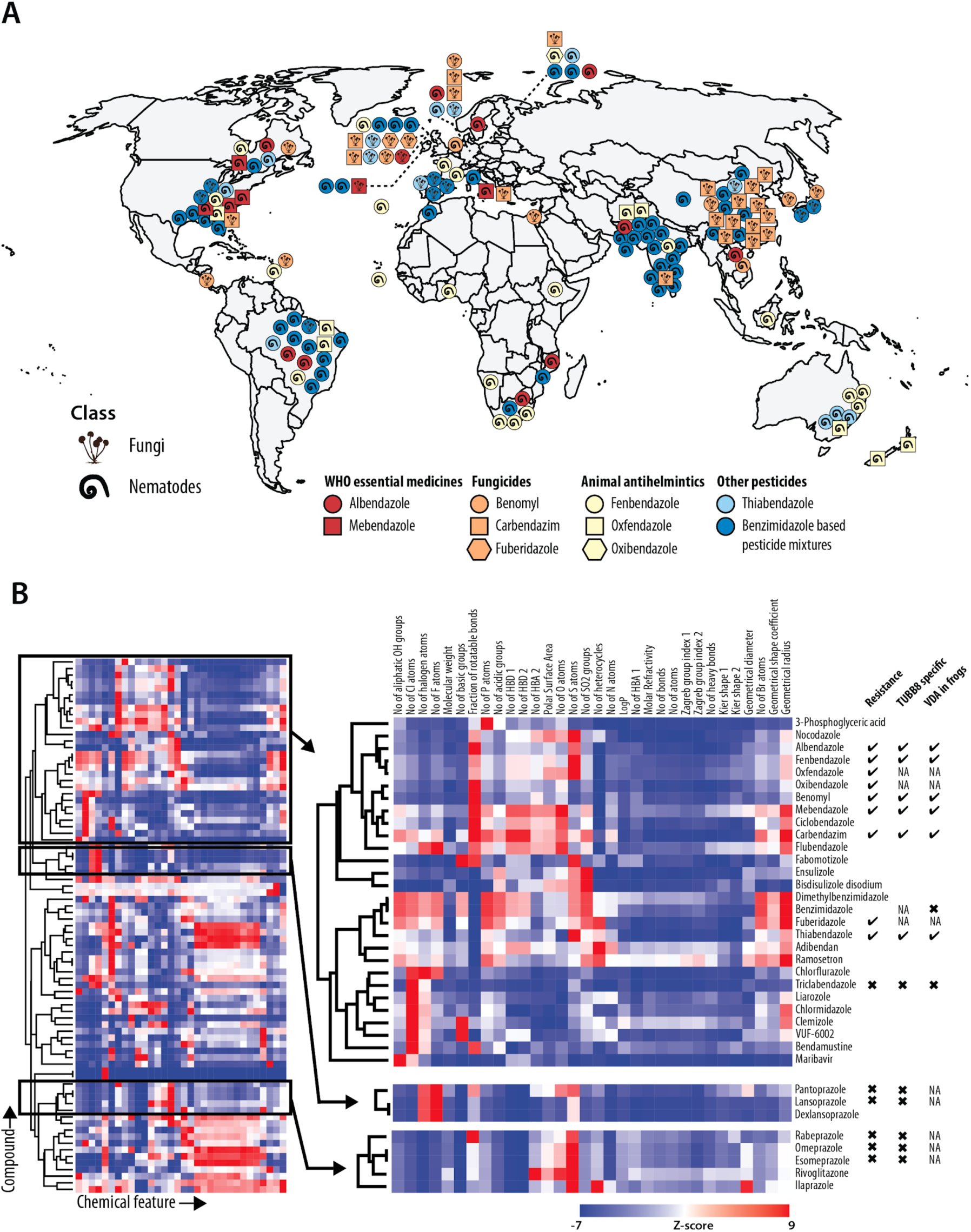
Global trends in benzimidazole resistance mutations and chemical structural similarities suggest numerous potential vascular disrupting agents. (**A**) 3 β-tubulin mutations, (F167Y, E198A, F200Y) conferring benzimidazole resistance have been globally observed among parasitic nematode and fungal species. Each icon represents an instance of β-tubulin suppressor mutations occurring in benzimidazole resistant parasitic fungal or nematode species (See **File S1** for list of species showing benzimidazole resistance). (**B**) Commonly used benzimidazoles hierarchically clustered by their chemical properties suggest new vascular disrupting agents with similar molecular mechanisms to TBZ. (Left) Clustergram of 81 widely used benzimidazole compounds spanning a wide range of drug classes grouped by chemical features (See **File S5** for the full list of compounds and features analysed). (Right) Zooms of black boxes indicate 3 clades containing TUBB8 specific VDA candidates (top) and proton-pump inhibitors (bottom).

In order to understand how extensively distributed benzimidazole resistance was, we mined ∼40 years of literature to identify reported cases of pesticide resistant species seen in wild and parasitic nematodes and fungi. Benzimidazole resistance is a global phenomenon (**Fig. 4A**); across 9 major commercial benzimidazole-based pesticides, we found multiple independent instances of reported resistance across 27 (12 nematodes and 15 fungal) parasitic species (**File S1**), all of which exhibited at least 1 of the 3 signature β-tubulin mutations. These widespread patterns of benzimidazole pesticide resistance suggested at least 9 new candidate VDAs.

As a complement to the epidemiological data, we also considered chemical properties by asking if pesticide benzimidazoles shared similar chemical feature profiles relative to other benzimidazoles. We curated >80 commercially available compounds in the benzimidazole class spanning a diverse range including pesticides, fungicides, therapeutics, and preservatives. Upon hierarchical clustering of these benzimidazoles based on their chemical properties computed from JOELib’s features matrix^72,73^ (**File S5**), we found that pesticide benzimidazoles generally shared similar chemical properties and clustered together (**Fig. 4B**).

### Numerous commercially used benzimidazoles also function as vascular disrupting agents

We next tested if pesticides exhibiting the epidemiological signature and clustering in the same clades by virtue of their chemical features would also specifically inhibit TUBB8 and function as VDAs. We selected 12 commercially used benzimidazole compounds across 2 clusters (**Fig. 4B**). Our list included 2 anthelmintics, both World Health Organization essential medicines (albendazole and mebendazole) prescribed to treat broad-spectrum human intestinal nematode infections; fenbendazole, an anthelmintic prescribed specifically for animals against gastrointestinal nematode parasites; 2 currently banned pesticides, benomyl and carbendazim, formerly used in agriculture; triclabendazole, specifically used to treat liver fluke infections; and 5 proton-pump inhibitors (esomeprazole, lansoprazole, omeprazole, pantoprazole, and rabeprazole) used to treat gastrointestinal and stomach acid disorders. The latter set were from a different clade and did not exhibit the epidemiological signature, serving as negative controls.

We first took advantage of our humanized yeast strains to rapidly discriminate TUBB8-specific inhibition from general β-tubulin inhibition. We found that 5 of the 12 compounds tested selectively inhibited TUBB8, as evidenced by the growth profiles observed for the humanized strains when cultured in the presence of the drugs (**Fig. 5, S4**). Notably, none of the 5 proton pump inhibitors or colchicine exhibited any tubulin inhibition (**Fig. S4, S5A**), confirming the specificity of the epidemiological signature as a predictor of TUBB8 inhibition. In contrast, triclabendazole was generally toxic, behaving as a pan-isotype inhibitor (**Fig. S5B**).

**Figure 5.**
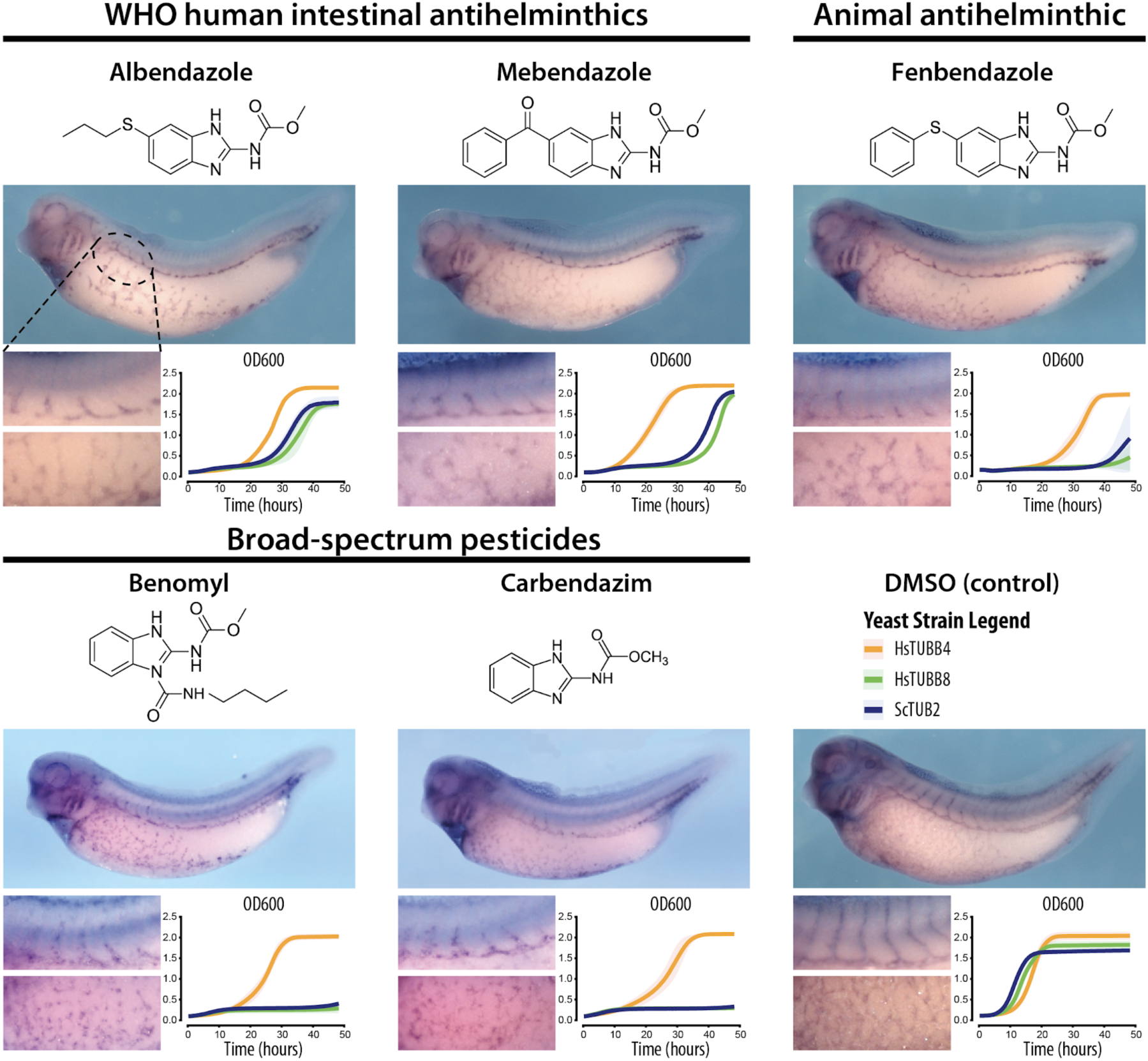
Commercially used benzimidazole pesticides, antifungals, and antihelminthics are also TUBB8-specific and disrupt vasculature. *In situ* hybridization of blood vessels (using the *erg/flk1* probe^18^) in *Xenopus laevis* embryos indicate the disruption of the vasculature caused by the presence of human and animal antihelminthics and broad-spectrum pesticides as compared to the DMSO control. Insets show growth profiles for yeast strains with humanized β-tubulin TUBB4 (orange) and TUBB8 (green) compared to wild-type (blue) when grown in the presence of each compound.

Testing the 5 positive TUBB8-inhibiting compounds in *Xenopus laevis* embryos showed strong vascular disrupting activity for all 5 compounds (**Fig. 5**). As we observed previously for TBZ^20^, the gross morphology of the treated embryos was largely normal (**Fig. 5)**. Thus, this broader class of benzimidazoles do in fact generally act as vascular disrupting agents in vertebrates.

## DISCUSSION

In the >30 years of therapeutic research efforts in the angiogenesis field only a highly restricted set of drugs have yet been approved^11^. Given the frequent failure to successfully make it through clinical trials and the high costs and lengthy process associated with developing new compounds, drug repurposing can offer efficient alternatives in developing new patient therapies with accelerated timeframes. This study represents a rather unconventional path to drug repurposing, leveraging a combination of model organisms, humanized yeast, cell culture, molecular modeling, and epidemiological data mining to determine TBZ’s molecular target and mechanism of vascular action. Indeed, TBZ was initially identified as a VDA and angiogenesis inhibitor by using a Baker’s yeast model of angiogenesis discovered in a computational search for orthologous phenotypes, or phenologs, aimed at exploiting deep evolutionary conservation to prioritize yeast processes relevant to human diseases^18–20^. Although obviously lacking blood vessels and a circulatory system, yeast nonetheless retains conserved biological pathways and processes relevant to vertebrate angiogenesis genes, and it was on the basis of these conserved processes that the antifungal compound TBZ was initially suspected, later confirmed, to be an angiogenesis inhibitor^20^.

While TBZ somewhat reduced the abundance of tubulin proteins in human cells^20^, at angiogenesis-inhibiting doses, the overall morphology of TBZ treated animals was normal, suggesting that only certain cell types, specifically those endothelial cells involved in forming the vasculature, might be uniquely susceptible to TBZ. Here, we find that TBZ does indeed specifically modulate the microtubules in vascular endothelial cells. Several currently identified microtubule targeting drugs have been reported to interfere with polymerization dynamics by binding β-tubulin^11,16^ close to or at the colchicine binding site. Building on previous work^27,48^, our *in silico* modeling results suggest that TBZ’s binding site, while in close proximity to the colchicine binding site, is distinct from it, thereby uncovering a novel β-tubulin effector site likely specific to other benzimidazoles and TBZ analogs.

In contrast to β-tubulin anticancer drugs, which have largely shown pan-isotype activity, to our knowledge, this study presents an unusual case of isotype-specific drug targeting in the β-tubulin gene family. Fungal suppressor studies on benzimidazole resistance have repeatedly found resistant mutations in β-tubulin; we found that 8 of 9 human β-tubulins natively harbor the same suppressor mutations and consequently exhibit unfavourable steric clashes interfering with TBZ binding. We demonstrate both *via* human cell culture microtubule assays and humanized yeast drug sensitivity tests that TBZ selectively targets only TUBB8 among the nine human β-tubulins, thus disrupting microtubule dynamics and reducing end-binding protein accumulation at the plus ends of microtubules in susceptible cells.

With TUBB8 thus acting as the specific target, it follows that of all human cell types, vascular endothelial cells must in turn be particularly sensitive to inhibition of TUBB8, leading to selective disruption of the vasculature relative to other human tissues. It remains to be seen why TBZ’s vascular disrupting activity is restricted to immature or newly forming blood vessels, but we speculate that this subset of the vasculature lacks reinforcing cell-cell contacts typical of larger, more established vasculature, leading to greater sensitivity to TBZ-induced microtubule disruption. As β-tubulin isotypes tend to be broadly expressed and often substitute for one another in microtubule structures^74,75^, one possibility is that TUBB8 inhibition simply leads to the loss of interactions with endothelial cell-specific components, thus specifically impacting vasculogenesis/angiogenesis. However, gene-gene and gene-drug interactions can often proceed by less obviously direct mechanisms to selectively impact cell types or phenotype penetrance *via* conditional cell-specific or dosage-dependent synthetic interactions^76,77^. It would thus not be surprising for the consequences of inhibiting TUBB8 in vascular endothelial cells to be similarly indirectly mediated by endothelial cell-specific synthetic interactions. Further experiments characterizing TBZ’s selective activity against newly forming/formed vasculature and the vascular-specific roles of TUBB8 in tetrapods could offer valuable insights into the cytoskeletal dynamics underlying vasculogenesis and angiogenesis.

Based on chemical properties and signature resistance mutations observed against benzimidazole compounds, we identified a larger class of extensively used fungicides and pesticides that all exhibit vascular disruption activity. While our results suggest possible new clinical applications for these compounds, they also highlight the potential caveats of their use in at-risk populations, especially for the two compounds (albendazole and mebendazole) that are FDA approved for human use. The WHO recommends the use of both albendazole and mebendazole as essential antihelmenthics worldwide for children up to the age of 14 against soil-transmitted helminth infections. Moreover, these compounds are widely used as public health interventions in pregnant women after the first trimester in regions where hookworm and whipworm infections exceed 20%^78,79^. While in the US, the risk of mebendazole use during pregnancy has not been assigned, our data add weight to WHO recommendations that these drugs should not be administered in the first trimester of pregnancy and suggest their use be carefully evaluated in patients in which angiogenesis inhibition might pose risks, including using caution later in pregnancy in light of the evidence that the compounds disrupt immature vasculature and might prove harmful to a developing fetus. Conversely, while efforts in the angiogenesis field have been often motivated towards developing anticancer therapies, the wide use of the compounds discussed here and their FDA-approved status could open alternative paths to treating other angiogenesis and/or vascular related diseases, such as diabetic retinopathy, macular degeneration, and hemangioma. It remains to be seen if other benzimidazoles sharing similar chemical profiles to those tested in our work (such as ciclobendazole, nocodazole, oxibendazole, and oxfendazole) also exhibit vascular disrupting activity.

More broadly, our framework of leveraging phenotypic relationships between species and repurposing model organisms to systematically explore drug mechanisms opens new routes for drug repurposing and discovery, and highlights the power of systems biology and evolution-guided approaches in advancing our knowledge of conserved genetic modules and how their disruption manifests in disease. This work also illustrates how duplicated genes diversify their functions and reinforces the therapeutic benefits of finding drugs specific to individual gene family members. As evidenced by the high degree of replaceability of conserved genes from cross-species complementation assays^80–82^, we anticipate that the combination of humanized yeast and phenolog-based disease modeling can be extended beyond vascular disruption to other conserved processes and therapies targeting them.

## MATERIALS AND METHODS

### Multiple sequence alignment

Human gene sequences were downloaded from the Uniprot database. The multiple sequence alignment for *S. pombe, S. cerevisiae*, and 9 human β-tubulin genes was constructed using MAFFT v7^83^ and visualized in Geneious v10 (https://www.geneious.com).

### Molecular modeling of β-tubulins

Homology models of human and fungal β-tubulins were constructed using as a reference structure the previously determined *Ovis aries* β-tubulin crystal structures (PDB: 3UT5 and 3N2G)^46,47^. The template was prepared using the Molecular Operating Environment (MOE.09.2014) software package from Chemical Computing Group. The structure was inspected for anomalies and protonated/charged with the Protonate3D subroutine (310K, pH 7.4, 0.1 M salt)^84^. The protonated structure was then lightly tethered to reduce significant deviation from the empirically determined coordinates and minimized using the Amber10:EHT forcefield with R-field treatment of electrostatics to an RMS gradient of 0.1 kcal mol^-1^ Å^-1^. Homology models of the wild-type fungal β-tubulin were prepared by creating 25 main chain models with 25 sidechain samples at 298K (625 total) within MOE. Intermediates were refined to an RMS gradient of 1 kcal mol^-1^ Å^-1^, scored with the GB/VI methodology, minimized again to an RMS gradient of 0.5 kcal mol^-1^ Å^-1^ and protonated. The final model for each variant was further refined by placing the protein within a 6 Å water sphere and minimizing the solvent enclosed structure to an RMS gradient of 0.001 kcal mol^-1^ Å^-1^. Models were evaluated by calculating Phi-Psi angles and superimposed against the reference structure. Homology models for each human β-tubulin were prepared similarly, based on generating a total of 625 models and averaging to make a final model for each β-tubulin isotype.

### *In silico* docking of TBZ into β-tubulins

Potential binding sites were evaluated using the Site Finder application and recent computational work on benzimidazole binding to parasitic β-tubulins^27,85^. Conformational variants of TBZ were created in 3-D within MOE. A database of conformations was then used to dock TBZ to the wild-type homology model using induced fit and template similarity protocols. The placement was scored with Triangle Matcher and rescored with London dG. Poses were refined with the Amber10:EHT forcefield with GVBI/WSA dG scoring. Candidate poses were then identified by inspecting polar contacts. Geometry optimization was carried out with MOPAC 7.0 using AM1. Conformational analysis of the bound structure was evaluated with LowModeMD^86^. 2-D contact maps were created using Ligand Interactions^87^.

### Cell culture

HUVEC cells were purchased from Clonetics and were used between passages 4 and 9. HUVECs were cultured on 0.1% gelatin-coated (Sigma) plates in endothelial growth medium-2 (EGM-2; Clonetics) in tissue culture flasks at 37 °C in a humidified atmosphere of 5% CO2. NIH-3T3 cells were obtained from Vishy Iyer at the University of Texas at Austin and cultured in Dulbecco’s Modified Eagle’s Medium (DMEM) with 10% bovine calf serum.

### Immunohistochemistry

Cell lines were cultured in 6-well plates and treated with thiabendazole dissolved in 1% DMSO. Control cells received 1% DMSO. After 24 h, cells were fixed with methanol at −20 °C for 10 min and subsequently with 4% paraformaldehyde in PBS at room temperature for 10 min. Cell membranes were permeabilized with 0.2% Triton X-100 in PBS, and nonspecific antibody binding sites were blocked with 5% goat serum for 1 h at room temperature. Cells were incubated with primary antibodies to EB1 (BD Bioscience) and α-tubulin (Sigma) at 4 °C overnight. After washing with PBST, primary antibodies were detected by Alexa Fluor-488 or 555 goat anti-rabbit or mouse immunoglobulin (IgG). 4′,6-Diamidino-2-phenylindole (DAPI dye, Sigma) was added as needed to visualize nuclei.

### Cell transfection and perfusion

EB3-eGFP cDNA obtained from Anna Akhmanova was cloned into the vector CS2+^88^. TUBB4 (Origene, RG203945) and TUBB8 (Origene, RG213889) cDNAs were purchased and cloned into the vector CS107-RFP-3Stop. HUVEC cells were transfected by nucleofection (Lonza) according to the manufacturer’s instructions. To analyze the effect of TBZ in living cells, we used a closed perfusion system (POC-R2, Pecon) connected to a peristaltic pump (Ismatec). 1% DMSO, 250 µM TBZ or 1% DMSO diluted in EBM-2 medium was flowed at 100 µl/min rate for the indicated times.

### Western blotting

HUVECs were cultured in 6-well plates and treated with 1% DMSO or 1% DMSO, 250 µM TBZ for 24 hours. Cells were lysed in cell lysis buffer (Cell Signaling Technology) containing 1 mM PMSF and analyzed by SDS-PAGE and western blotting using anti-EB1 (BD Bioscience) or anti-EB3 (Millipore) or anti-Clip170 (Santa Cruz) antibodies.

### Imaging and image analysis

Immunohistochemistry experiments, live HUVECs, and live *KDR:GFP* transgenic *Xenopus laevis* were imaged using an inverted Zeiss LSM5 Pascal and Zeiss LSM700 confocal microscope, and super-resolution structured illumination (SR-SIM) combined with Zeiss LSM710 microscope. Comet lengths were measured using the software Fiji. Confocal images were cropped and enhanced in Adobe Illustrator and Adobe Photoshop for the compilation of figures.

### Benzimidazole clustering analysis

81 commercially used benzimidazole compounds spanning a wide range of classes were curated from PubChem^89^. JOElib (http://joelib.sourceforge.net), OpenBabel^90^, and Chem Mine features were computed using ChemMine tools^73^. Heatmaps were visualized using Morpheus (https://software.broadinstitute.org/morpheus). Clustergrams were generated by hierarchical clustering on the one minus Pearson correlation coefficient with average linkage.

### Humanizing yeast β-tubulin using CRISPR-Cas9

The human TUBB4 and TUBB8 open reading frames were integrated chromosomally (from start to stop codon) into *Saccharomyces cerevisiae* in place of the endogenous *TUB2* open reading frame using CRISPR/Cas9 genome editing as described in Akhmetov *et al*.^91^. Two sgRNAs were designed targeting the yeast *TUB2* locus using the Geneious (v10.2.6) CRISPR-Cas9 tools suite, purchased as oligos from IDT, and cloned into yeast CRISPR-K/O vectors using the yeast toolkit (YTK)^92^ to express a synthetic guide RNA sequence, Cas9 nuclease, and a selectable marker (URA3)^52,93^. Repair templates were constructed by PCR amplification of the human β-tubulin ORF (from the human ORFeome^94^) flanked by 75 bp of target chromosomal boundary at the *TUB2* locus to facilitate recombination *via* homology directed repair. BY4741 (S288C) yeast strains were co-transformed with the CRISPR/Cas9 vector and repair template using Zymo Research Frozen-EZ Yeast Transformation II Kit. Transformants were selected on SC-URA media. Surviving colonies were screened by colony PCR, and Sanger sequenced to confirm replacement.

### Humanized yeast growth assays

Assayed benzimidazole compounds were all dissolved in 100% DMSO to prepare stock solutions of 5 or 10 mg/ml based on solubility. Candidate VDA compounds were titrated in ranges of 5-1000 µg/ml into growth medium depending on solubility (**Fig. S5** lists specific concentrations) for subsequent growth assays. Liquid growth assays were performed in triplicate in 96-well format using a Biotek Synergy HT incubating spectrophotometer. Humanized tubulin strains were pre-cultured to saturation in YPD and diluted into 150 µL of media to have 0.05-0.1 × 10^7^ cells/ml. Assays were typically run for 48 hrs with absorbance measured every 15 min.

### *Xenopus* embryo manipulations and VDA assays

*Xenopus* embryos were reared in 1/3× Marc’s modified Ringer’s (MMR) solution. Each drug was treated to embryos from stage 31 until stage 38 with 10 µg/ml or 20 µg/ml in 1% DMSO diluted in 1/3X MMR. Embryos were fixed at stage 38 with MEMFA, and whole-mount *in situ* hybridization for *erg* was performed as described in Sive *et al*.^87^.

## ACKNOWLEDGMENTS

The authors thank Andrew Ellington for critical feedback and discussions. This research was funded by the American Heart Association Predoctoral fellowship (#18PRE34060258) to R.K.G., Army Research Office (W911NF-12-1-0390) to J.D.G., Natural Sciences and Engineering Research Council (NSERC) of Canada Discovery grant (RGPIN-2018-05089), CRC Tier 2 (NSERC/CRSNG-950-231904), and the Canada Foundation for Innovation and Québec Ministère de l’Économie, de la Science et de l’Innovation (#37415) to A.H.K., the National Institute of Child Health and Human Development (R01HD099191) to J.B.W., and from the Welch Foundation (F-1515) and National Institutes of Health (R35 GM122480) to E.M.M..

## AUTHOR CONTRIBUTIONS

Conceptualization and methodology, R.K.G., H.J.C., J.D.G., A.H.K., J.B.W., E.M.M.; Computational analyses, R.K.G., H.J.C., J.D.G.; Investigation, R.K.G., H.J.C, C.L., J.D.G.; Formal analysis and visualization, R.K.G., H.J.C., E.M.M.; Writing – R.K.G., H.J.C, J.B.W., E.M.M.

## Supplementary Figures

**Figure S1.**
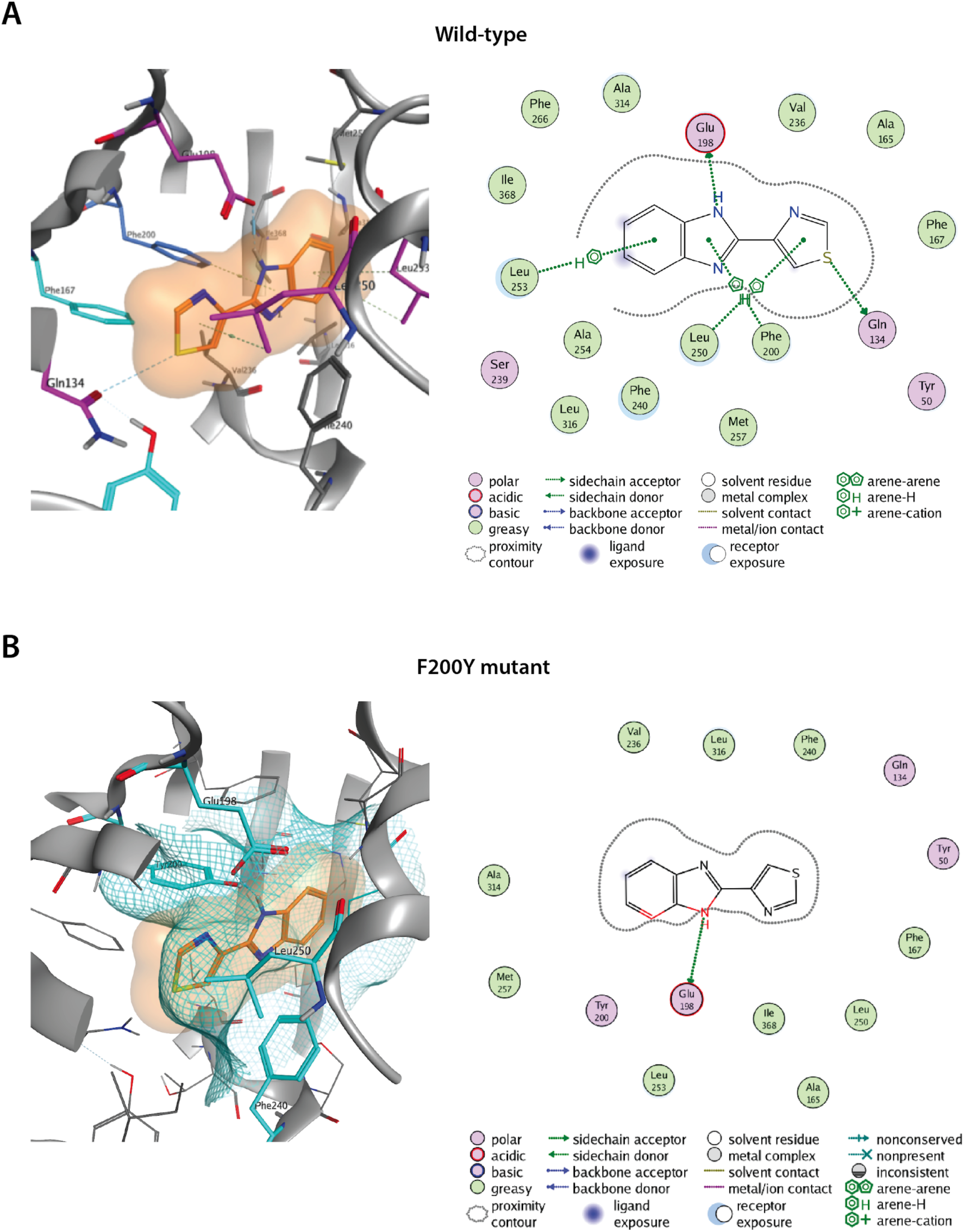
Homology modeling and *in silico* docking studies predict the TBZ binding site in the fungal β-tubulin *NDA3* structure. TBZ is well accommodated in wild-type *NDA3*’s predicted binding pocket (**A**) as opposed to its F200Y mutant (**B**). 3D structures and 2D contact maps shown on the left and right respectively indicate the steric clashes TBZ faces in the F200Y binding pocket. Cyan meshes (in 3D structures) and red highlights on the ligand (in 2D contact maps) indicate steric clashes in the binding pocket

**Figure S2.**
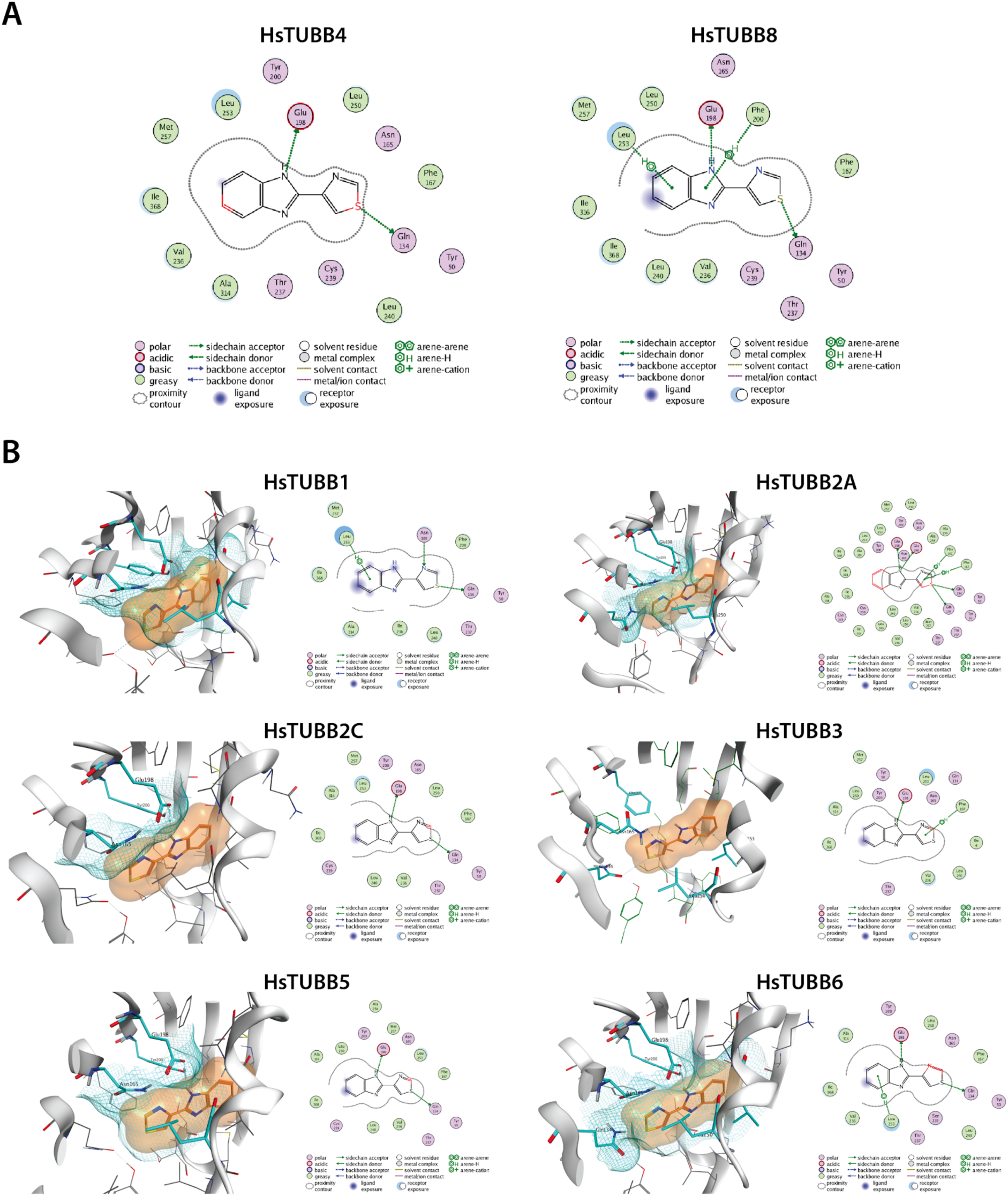
Only TUBB8 favorably binds TBZ among the 9 human β-tubulins. (**A**) 2D contact maps highlight ligand interactions between TBZ and TUBB4 (left) or TUBB8 (right). TBZ forms polar contacts with residues Q134, E198, F200, and L253 in TUBB8. In TUBB4, reorientation of the proposed binding pocket is observed. Substantial steric clashing is shown in red on TBZ. (**B**) *In silico* docking of TBZ into other human β-tubulin homology models suggests substantial steric clashes due to unfavorable binding pockets among 8/9 human β-tubulin isotypes. Cyan meshes (in 3D structures) and red highlights on the ligand (in 2D contact maps) indicate steric clashes in the binding pocket

**Figure S3.**
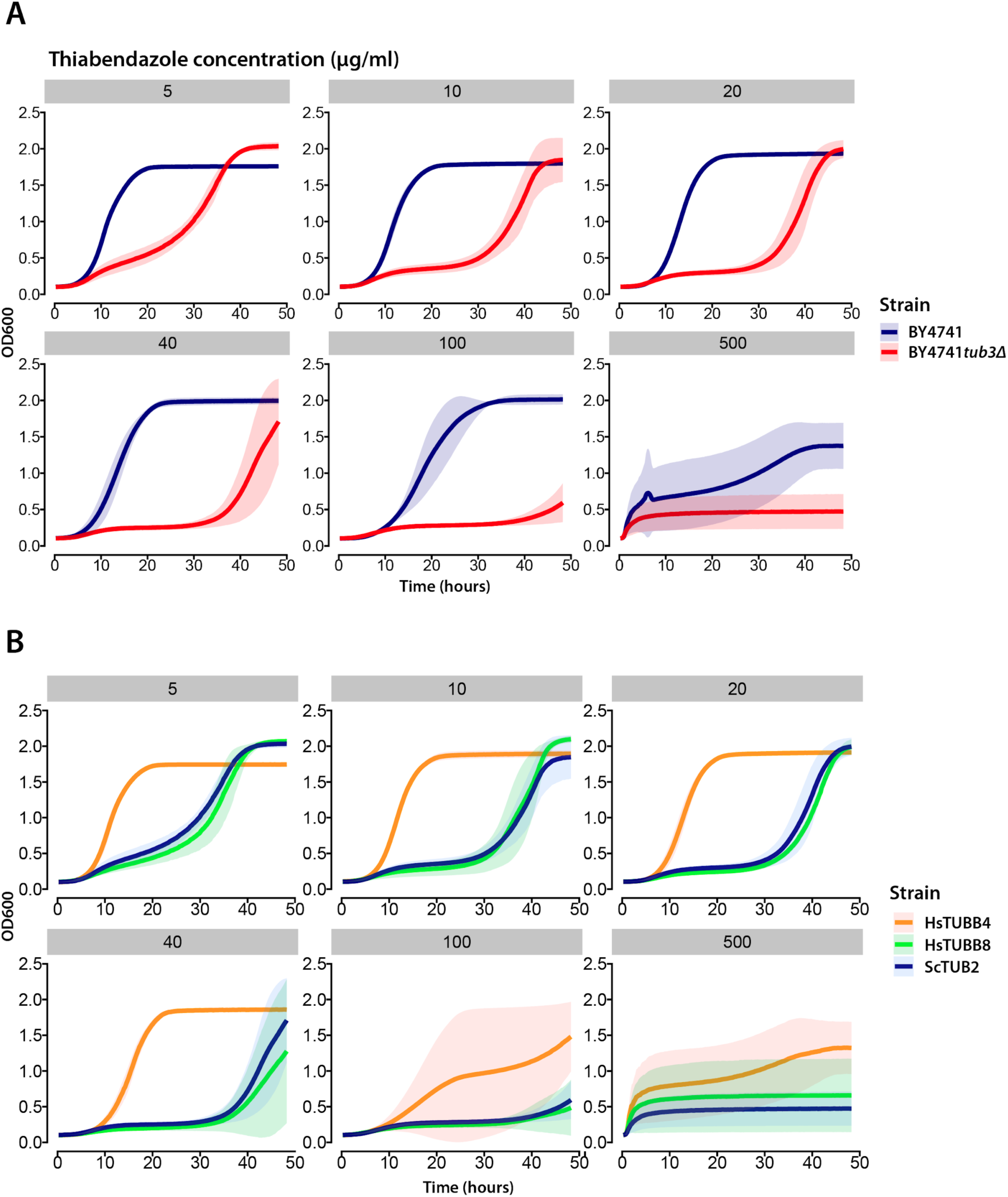
Yeast strains with modified β-tubulin are differentially sensitive to TBZ. (**A**) Deletion of yeast α-tubulin *TUB3* makes *Saccharomyces cerevisiae* (Baker’s yeast) sensitive to TBZ. (**B**) Growth profiles of *tub3Δ* yeast strains with wild-type *TUB2* (blue), humanized β-tubulins TUBB4 (orange), or TUBB8 (green) in increasing concentrations of TBZ show differential sensitivities to the drug.

**Figure S4.**
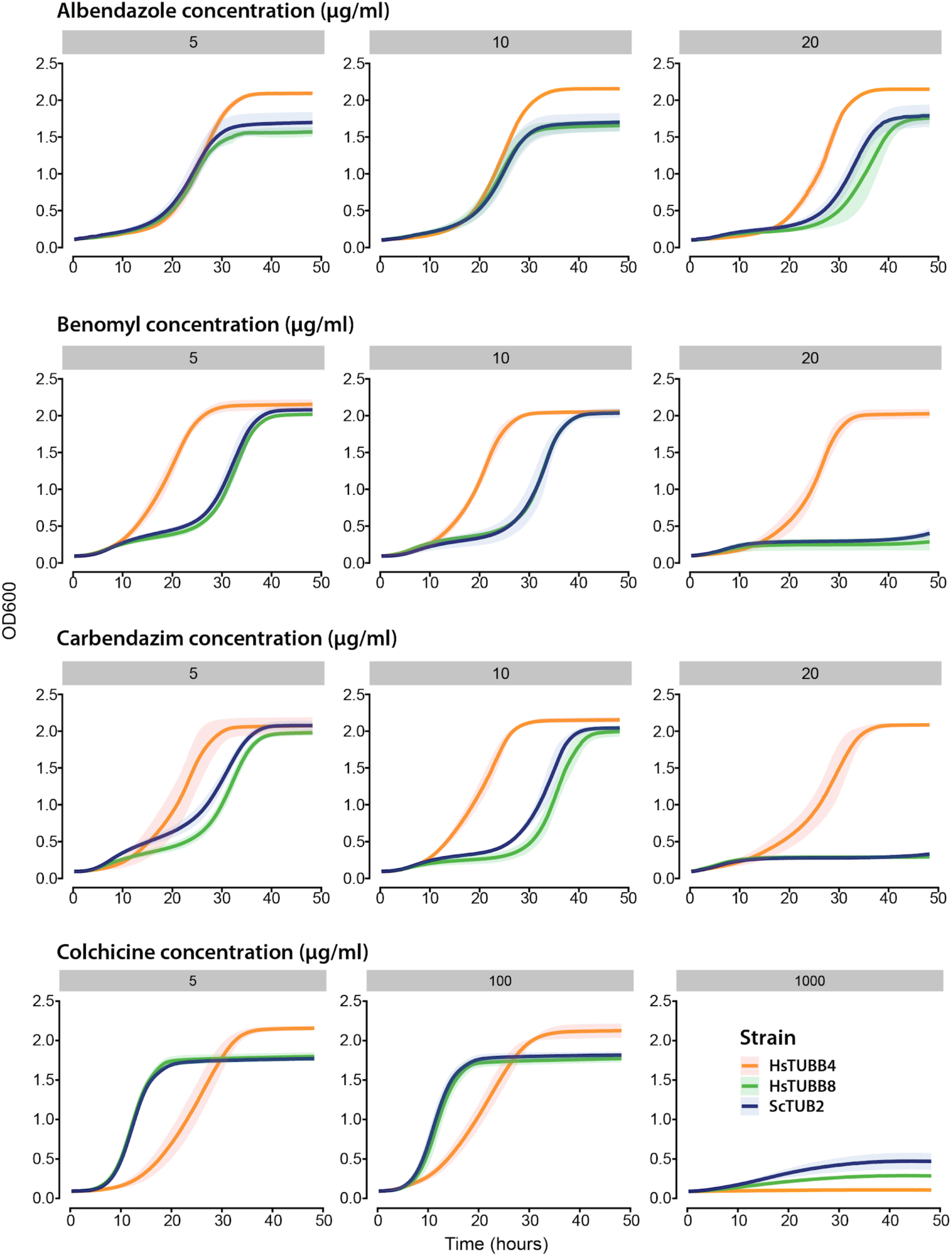
Growth profiles of benzimidazole treated yeast strains. Plots depict varying doses of albendazole, benomyl, carbendazim, and colchicine. Our data show that colchicine is a pan-isotype inhibitor whereas albendazole, benomyl, and carbendazim show some degree of specificity for TUBB8.

**Figure S5.**
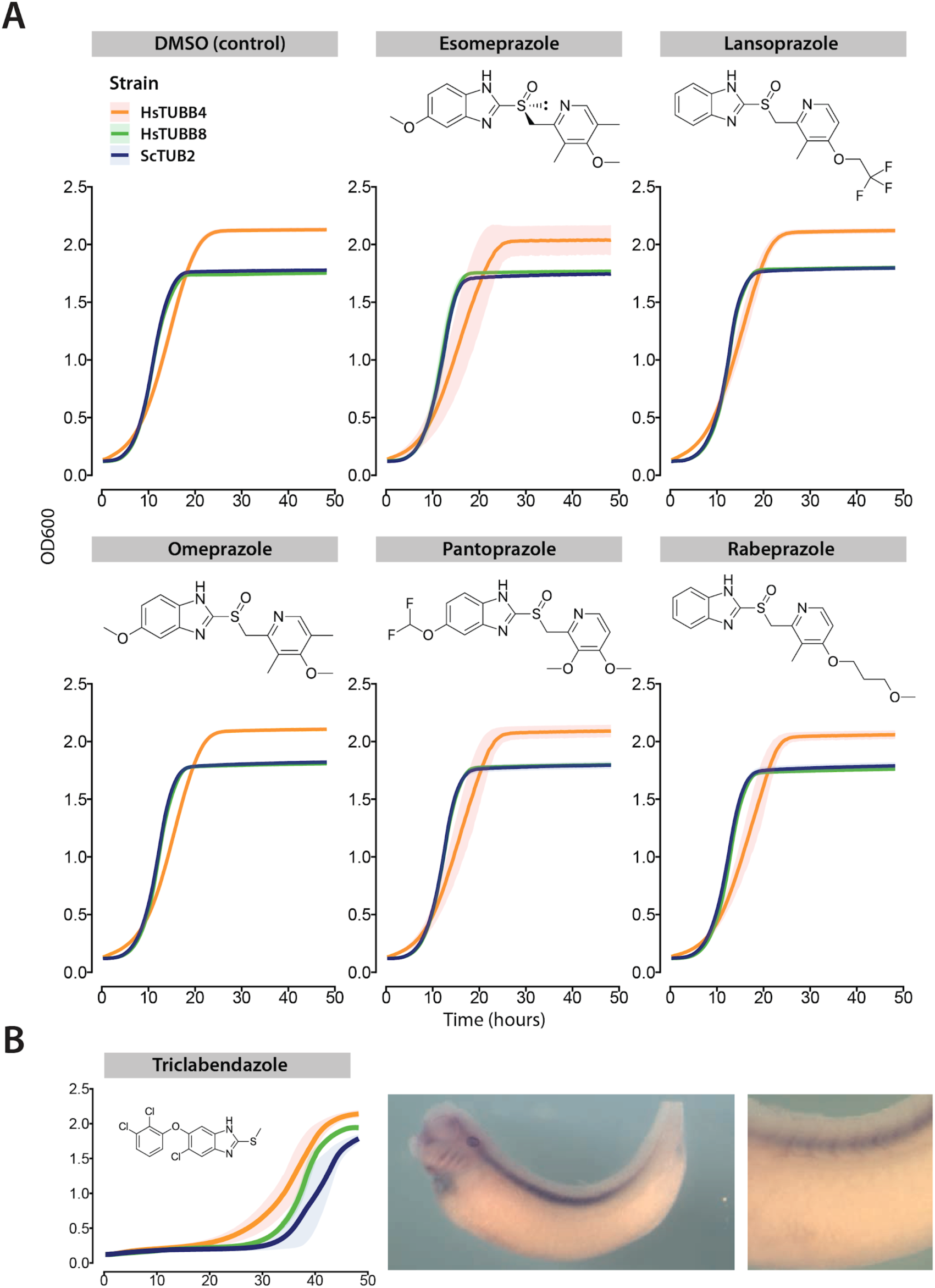
Proton-pump inhibitors do not elicit growth defects in humanized strains. (**A**) Yeast strains harboring the beta-tubulin gene *TUB2* (blue), TUBB4(orange), and TUBB8 (green) are not inhibited by proton pump inhibitors (drug conc. 40 µg/ml). (**B**) Triclabendazole is a pan-isotype β-tubulin inhibitor inhibiting both wild-type and humanized yeast strains (Left) and lethal to developing *Xenopus laevis* embryos at stage 38. (Right).

## Supplementary information

**File S1**. Species resistance table curated from literature

**File S2**. Yeast site finder statistics

**File S3**. Docking free energy scores across β-tubulin isotypes

**File S4**. Yeast wt β-tubulin induced fit TBZ docking scores

**File S5**. Benzimidazole chemical features

